# The genome of the reef-building coral *Porites harrisoni* from the southern Persian/Arabian Gulf

**DOI:** 10.64898/2026.02.26.708201

**Authors:** Anna Fiesinger, Abdoallah Sharaf, Rachel Alderdice, Gabriela Perna, Hannah Manns, John A. Burt, Christian R. Voolstra

**Affiliations:** Department of Biology, University of Konstanz, Konstanz, Germany; SequAna Core Facility, Department of Biology, University of Konstanz, Konstanz, Germany; Center for Genomics and Systems Biology (CGSB) and Mubadala Arabian Center for Climate and Environmental Sciences (Mubadala ACCESS), New York University Abu Dhabi, Abu Dhabi, United Arab Emirates

**Keywords:** coral thermal resilience, genome assembly, genome annotation, mitochondrial genome

## Abstract

We present a genome assembly from the coral species *Porites harrisoni* from the southern Persian/Arabian Gulf, the hottest ocean basin where corals live. The assembly is 626.7 Mb in size, spanning 1,883 contigs with a contig N50 of 807.4 kb, including a single-contig mitochondrial genome. The assembly has a BUSCO completeness of 86.3% (single = 72.5%, duplicated = 13.7%, fragmented = 1.2%, missing = 12.5%) using the eukaryota_odb10 reference set (*n* = 255). A total of 59.23% of the nuclear genome consists of repeats, comprising 15.89% retroelements, 10.00% DNA transposons, and 31.71% unclassified repeats. Gene annotation of this nuclear genome assembly identified 27,823 protein-coding genes. The mitogenome has an assembly size of 18,639 bp with 13 protein-coding genes as well as 2 tRNAs and 2 rRNAs. The genome of *P. harrisoni* provides a valuable genomic resource of a coral from an extreme environment, which will enable comparative analyses, enhancing our understanding of the genomic architecture underlying thermal resilience. Such comparisons will contribute to elucidating the evolutionary basis of heat tolerance and adaptive capacity of corals in the context of rapid climate change.

## Introduction

Corals surviving in extreme environments (i.e., experiencing variable and extreme water temperatures as well as high salinities) can serve as model organisms to study the adaptive capacity of corals in more temperate regions. Such corals offer valuable insight into the genomic architecture and genes underlying thermal resilience [1–3]. The hottest oceanic basin where corals occur is the shallow and semi-enclosed Persian/Arabian Gulf (PAG) where temperature extremes and ranges exceed those of coral reefs elsewhere (<12 °C to >36 °C annually) [4, 5]. Due to the relatively recent geological formation of the PAG, estimated to be about 6–12 ka old, and its shift to a hot climate, corals in these waters have had little time to adapt to these extreme conditions [6]. As a result, they inform our understanding of how coral reefs at large may respond to rapid warming.

One prominent coral in the PAG, and among the species most resilient to recurrent bleaching events [7], is the endemic species *Porites harrisoni* Veron, 2000 (Fig. 1) from the complex coral clade. The complex clade comprises many ecologically dominant reef builders [8, 9] with the genus *Porites* hosting several thermally resilient species [10–14]. Colonies of *P. harrisoni* abundantly inhabit shallow fringing reefs in the PAG [15], and *P. harrisoni* exhibits diverse growth forms – submassive, nodular, columnar, or branching – on a broad encrusting base, with coloration varying from light to dark brown [15]. The thermal tolerance of *P. harrisoni* is in part attributed to its association with the thermotolerant symbiotic alga *Cladocopium thermophilum*, especially in the southern PAG [6, 16, 17]. This alga has undergone rapid adaptive radiation upon colonizing the PAG [18]. In line with this, several studies suggest that PAG corals carry unique genomic signatures underpinning their exceptional heat tolerance [19, 20], which are heritable [20, 21]. Like many other corals in the PAG, *P. harrisoni* has one of the highest ecological bleaching thresholds in the world [22]. Originally described as a different species, *P. harrisoni* was first identified for its exceptional tolerance to extreme temperatures in a study highlighting the thermal tolerance of PAG corals [23] and later recognized for its persistence in hypersaline and thermally extreme lagoons where most corals are unable to survive [24]. However, it also lives exceptionally close to its upper thermal tolerance threshold [25]. Due to the increasing rate of ocean warming, this species may provide critical insight into the genomic basis of coral thermal resilience, as it has thus far managed to survive in harsh conditions where other corals, particularly in the southern PAG, have not as a result of recurrent bleaching [12].

**Figure 1.**
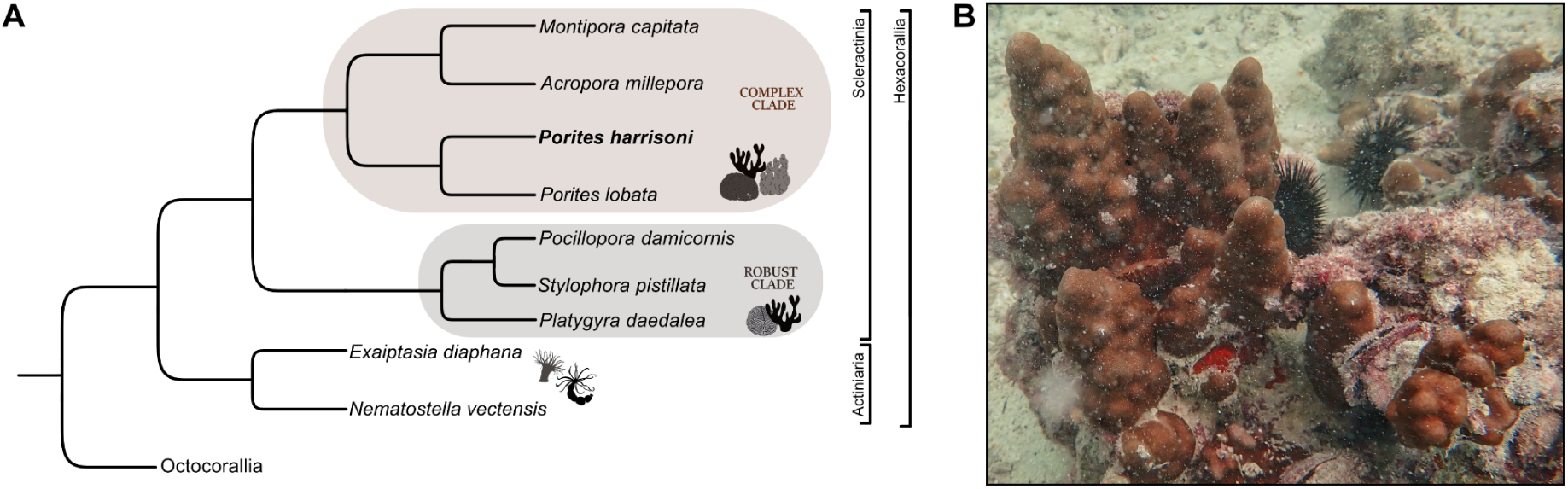
The species *Porites harrisoni*, Veron 2000. **(A)** Cladogram illustrating evolutionary relationships of *Porites harrisoni* (in bold) and other Hexacorallia with sequenced genomes [28–35]. The data is adapted from a simple species tree generated with the NCBI ETE v3.1.3 toolkit [36]. Robust and complex coral clades are indicated [8, 9, 37]. A full-scale phylogenetic tree illustrating evolutionary relationships of the genus *Porites* within the Scleractinia is available from a recent study [38]. **(B)** *In situ* colony photo of *P. harrisoni* from Saadiyat Reef in the Persian/Arabian Gulf. Picture credit: Christian R. Voolstra.

Here, we report on the sequencing, assembly, and annotation of the genome of *P. harrisoni* from the southern PAG. Species taxonomy for this species is Eukaryota; Opisthokonta; Metazoa; Eumetazoa; Cnidaria; Anthozoa; Hexacorallia; Scleractinia; Fungiina; Poritidae; *Porites*; *Porites harrisoni* Veron, 2000 (NCBI:txid627007). The genome assembly is based on long-read Oxford Nanopore Technologies (ONT) sequencing. The generated genome is 626.7 Mb in size, comprising 1,883 contigs with a contig N50 of 807.4 kb. The nuclear genome annotation yielded 27,823 protein-coding genes, which were validated through RNA-Seq and available curated Metazoa proteomes. In addition, we assembled a complete single-contig mitochondrial genome (mitogenome), which should inform biodiversity assessment and further contribute to resilience understanding [26, 27]. The genome of *P. harrisoni* will facilitate coral species comparisons regarding the genomic nature of thermal resilience.

## Material & Methods

### Sample collection, DNA isolation, and sequencing

One colony of *Porites harrisoni* was tagged and sampled for genomic DNA (gDNA) extraction for the reference genome (BioSample ID SAMN41390914; Table 1) at Al Saada Reef, Abu Dhabi, United Arab Emirates (SA; 24.085250, 52.243333) in the thermally extreme southern Persian/Arabian Gulf on 30 May 2022 using a Nemo underwater drill (Nemo Power Tools LLC, Nevada, USA) fitted with a diamond tip corer. The coral plug was sampled from a Coral Bleaching Automated Stress System (CBASS) [39] experiment described elsewhere [25] from the baseline temperature (34.1 °C) treatment at time point two (i.e., after overnight recovery). The coral fragment was submerged in 3 ml of DNA/RNA shield (Zymo Research Europe GmbH, Freiburg, Germany) in a WhirlPak bag for at least 3 hours, after which the tissue was airpicked off the skeleton using an air pump with sterile filter tips. Tissue slurry was transferred to 2 x 2 ml cryotubes and stored at 4 °C until shipment to the lab facilities at the University of Konstanz, Germany. The sample was stored at –20 °C until extraction.

**Table 1.**
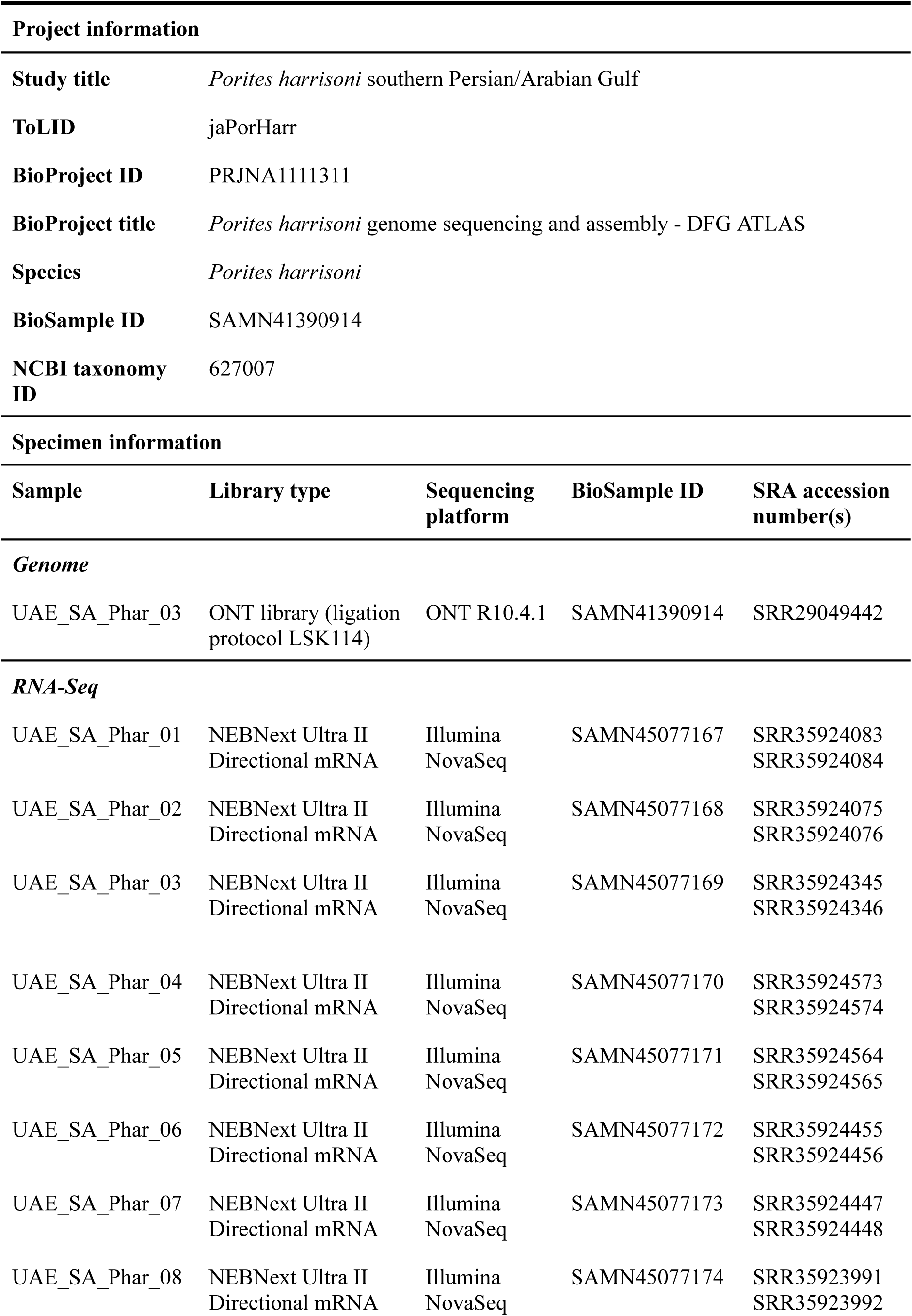
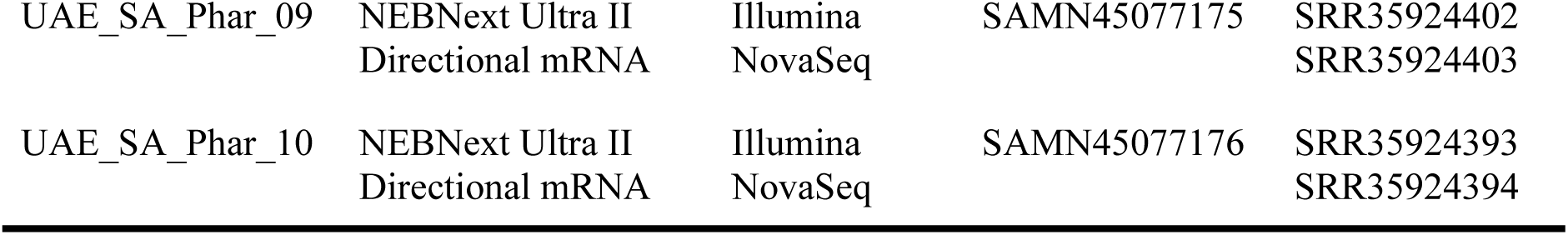
Specimen and sequencing data for *Porites harrisoni*.

DNA was extracted using the DNeasy Blood & Tissue kit (Qiagen, Hilden, Germany) following the manufacturer’s protocol with a minor modification. Briefly, the tissue slurry was homogenized using a Polytron PT 1200 E (Kinematica, Switzerland), the sample was spun down at 5,000 x *g* for 3 mins, and 90 µl of the supernatant were transferred from the cryotube to a 1.5 ml tube. Subsequently, 90 µl of ATL buffer and 20 µl of Proteinase K (Qiagen, Hilden, Germany) were added, and the sample was lysed for 1 hr at 56 °C at 300 rpm. After extraction, two elution steps were applied [40]: 50 µl of AE elution buffer (Qiagen, Hilden, Germany) were added to the membrane of the spin column, it was incubated for 1 min, and centrifuged at 7,000 x *g* for 1 min. A subsequent 50 µl AE buffer were added to the membrane of the spin column in the same elution tube, it was incubated for 5 mins and centrifuged as previously. The resulting DNA quality and quantity were assessed using a NanoDrop2000 spectrophotometer (Thermo Fisher Scientific, Waltham, Massachusetts, USA).

A total of 6.54 µg DNA were submitted to the NGS Competence Centre Tübingen (NCCT) for the construction of an ONT sequencing library using the ligation protocol LSK114 that was subsequently sequenced with an R10.4.1 flowcell on a PromethION 24 (Oxford Nanopore Technologies, Oxford, UK). Base calling was done using MinKNOW v23.07.12 (Oxford Nanopore Technologies, Oxford, UK) and Dorado v0.7.4. [41]. Base-called reads were trimmed using porechop v0.2.4 [42] and the quality was assessed using NanoPlot v1.44.1 [43]. For subsequent genome assembly, the ONT reads were split into longer reads for assembly (minimum average quality 3, minimum length 1000 bp) and shorter reads for polishing (minimum average quality 5, minimum length 500 bp), while removing reads mapping to algal symbionts (i.e., all publicly available reads from genomes of the dinoflagellate family Symbiodiniaceae) using chopper v0.9.0 [43]. The Symbiodiniaceae reference reads (NCBI:txid252141) were downloaded from NCBI using datasets v16.14.0 (parameters: genome taxon 252141) [44].

### Sample collection, RNA isolation, and sequencing

A total of 20 samples (∼2 cm each) were drilled from 10 colonies of *Porites harrisoni* at Al Saada Reef (SA; 24.085250, 52.243333) in the southern Persian/Arabian Gulf on 30 May 2022 for RNA isolation (Table 1) using a Nemo underwater drill (Nemo Power Tools LLC, Nevada, USA) fitted with a diamond tip corer. The coral plugs were sampled from a CBASS experiment [39] described elsewhere [25] from the baseline temperature (34.1 °C) treatment from time points one and two, i.e., after heat-hold and after overnight recovery. As described above, the coral fragments were submerged in 3 ml of DNA/RNA shield (Zymo Research Europe GmbH, Freiburg, Germany) in a WhirlPak bag for at least 3 hours, after which the tissue was airpicked off the skeleton using an air pump with sterile filter tips. Tissue slurry was transferred to 2 ml cryotubes and stored at 4 °C until shipment to the lab facilities at the University of Konstanz, Germany. The samples were stored at –20 °C until extraction.

RNA was extracted following a previously described protocol [45] using the RNeasy Mini Kit (Qiagen, Hilden, Germany) adapted for use with a QIAcube (Qiagen, Hilden, Germany). The quantity and quality of the RNA was assessed using a NanoDrop2000 spectrophotometer (Thermo Fisher Scientific, Waltham, Massachusetts, USA) as well as gel electrophoresis to check for the presence of bands. The extracted RNA was sent to the NCCT for library preparation using the NEBNext Ultra II Directional mRNA kit and subsequent paired-end sequencing on an Illumina NovaSeq (2 x 150 bp).

### Estimation of the genome size

The genome size was estimated using a k-mer histogram generated with Meryl v1.3 [46] based on the ONT base-called and trimmed reads, and visualized in GenomeScope2.0 [47].

### Genome assembly, curation, and evaluation

The initial draft assembly was done using a subset of size-selected ONT reads (minimum average quality 3, minimum length 1000 bp; total number of reads: 20,260,201) with the *de novo* Nanopore assembler NECAT v0.0.1 [48]. The quality of the assembled draft genome was assessed using BlobToolKit v4.3.0 [49], which prompted the removal of 442 contigs due to their similarity to non-related taxa. The draft assembly was then polished using a second subset of size-selected ONT reads (minimum average quality 5, minimum length 500 bp; total number of reads: 21,513,716) with one round of Racon v1.5.0 [50] and one round using Medaka v1.12.0 (parameters: --model r1041_e82_400bps_hac_v4.3.0) [51]. After polishing, funannotate v1.8.5 [52] was used to remove duplicated contigs and those shorter than 200 bp (funannotate clean; -m 200), prompting the removal of a further five contigs. The quality of the assembly was assessed using BUSCO v6.0.0 scores [53] and the BlobToolKit v4.3.0.

### Mitochondrial genome assembly and annotation

The mitogenome was assembled using the same set of size-selected reads as above (minimum average quality 3, minimum length 1000 bp), with a subsequent filtering step: reads were mapped to the available *Porites harrisoni* mitogenome (GenBank accession: MG754070, RefSeq accession: NC_037435) [54] using minimap2 v2.28 [55]. The mapped reads were then extracted using SAMtools v1.21 [56] and subsequently assembled using Canu v2.3 [57] (parameters: genomeSize=18k, -nanopore -trimmed). The resulting mitogenome was circularized using Circlator v1.5.5 (parameters: --assembler canu, --merge_min_id 85, --merge_breaklen 1000) [58]. The circularized mitogenome was then polished using Racon v1.5.0 with the shorter reads as above (minimum average quality 5, minimum length 500 bp). The mitogenome was annotated with MITOS2 [59], and manually curated in Geneious Prime v2025.1.1 (GraphPad Software, LLC, Boston, USA) against the published *P. harrisoni* mitogenome [54] and the mitogenome of the closely related species *Porites lobata* [60].

### Repeat annotation

DNA repeats in the genome were identified using a combination of RepeatModeler v2.0.4 [61] and the EDTA v2.2.2 pipeline [62], and masked using RepeatMasker v4.1.6 [63].

### Gene prediction and annotation

Gene prediction was performed using BRAKER3 v3.0.8 [64]. First, the transcript evidence was prepared as follows: After quality checking the RNA-Seq reads using FastQC v0.12.1 [65], the reads were trimmed using Trimmomatic v0.39 [66] and subsequently mapped to the *Porites harrisoni* assembly using STAR v2.7.11b [67]. The resulting mapping files were merged, and the reverse strand-specific RNA reads were extracted using SAMtools v1.21. Second, curated Metazoa proteomes were retrieved from OrthoDB 11 (https://bioinf.uni-greifswald.de/bioinf/partitioned_odb11/) and used as protein evidence as recommended by the BRAKER3 pipeline [64, 68, 69]. Lastly, BRAKER3 was run using the reverse strand-specific RNA mapping files and protein evidence as input with the masked genome assembly (parameters: --busco_lineage=metazoa_odb10, --species=Porites_harrisoni) [53, 70–79]. Using the script stringtie2utr.py from the BRAKER3 suite, untranslated regions (UTRs) were added to the annotation file output by BRAKER3. In addition, 674 eukaryotic high-confidence transfer RNAs were predicted using tRNAscan-SE v2.0.12 [80] based on filtering of the initial set of 5,938 putative tRNAs using EukHighConfidenceFilter. The high-confidence set of tRNAs was merged with the BRAKER3 structural predictions. Then, the annotation file was checked for overlapping genes using AGAT [81] and validated using GenomeTools v1.6.5 [82]. Finally, GffRead [76] was used to extract the predicted protein sequences from the merged file to use in the functional annotation. The predicted genes were annotated using InterProScan v5.59-91.0 [83], EggNOG-mapper v2.1.12 [84, 85], and Phobius v1.01 [86]. The respective annotation files were then fed into funannotate with the predicted genes in gff3 file format to synthesize all annotations (parameters: --species "Porites harrisoni" --busco_db metazoa, --eggnog --iprscan --phobius). The final annotation was assessed using BUSCO v6.0.0. The gene prediction and functional annotation outlined in this study were used in the development of the fully automated annotation pipeline GeneForge v1.0 [87].

## Results & Discussion

### Genome assembly

The genome was sequenced from a single *Porites harrisoni* colony, collected from Al Saada Reef (SA) in the southern Persian/Arabian Gulf on 30 May 2022 (Table 1). A total of 22,825,985 ONT long-reads were obtained from an R10.4.1 flow cell (Oxford Nanopore Technologies, Oxford, UK) following adapter removal and trimming, with a mean read length of 5,071 bp (median 3,407 bp) and a read length N50 of 7,424 bp. The reads showed a mean base call quality score of 14.9, and the longest read sequenced was 991,055 bp with a quality score of 10.

The estimated genome size using a k-mer approach (k = 21) was 599 Mb (Fig. 2). The analysis revealed a diploid genome (p = 2) that is predominantly homozygous (98% AA) with very little heterozygosity (1.96% AB). While GenomeScope profiles are not optimized for standard ONT reads (given the lower accuracy in comparison to short-reads), the use of ONT-derived k-mer counts still provides a valuable and informative approximation of genome size, particularly in the absence of complementary short-read sequencing. The genome size estimated here corresponds to other assembled *Porites* genomes [35].

**Figure 2.**
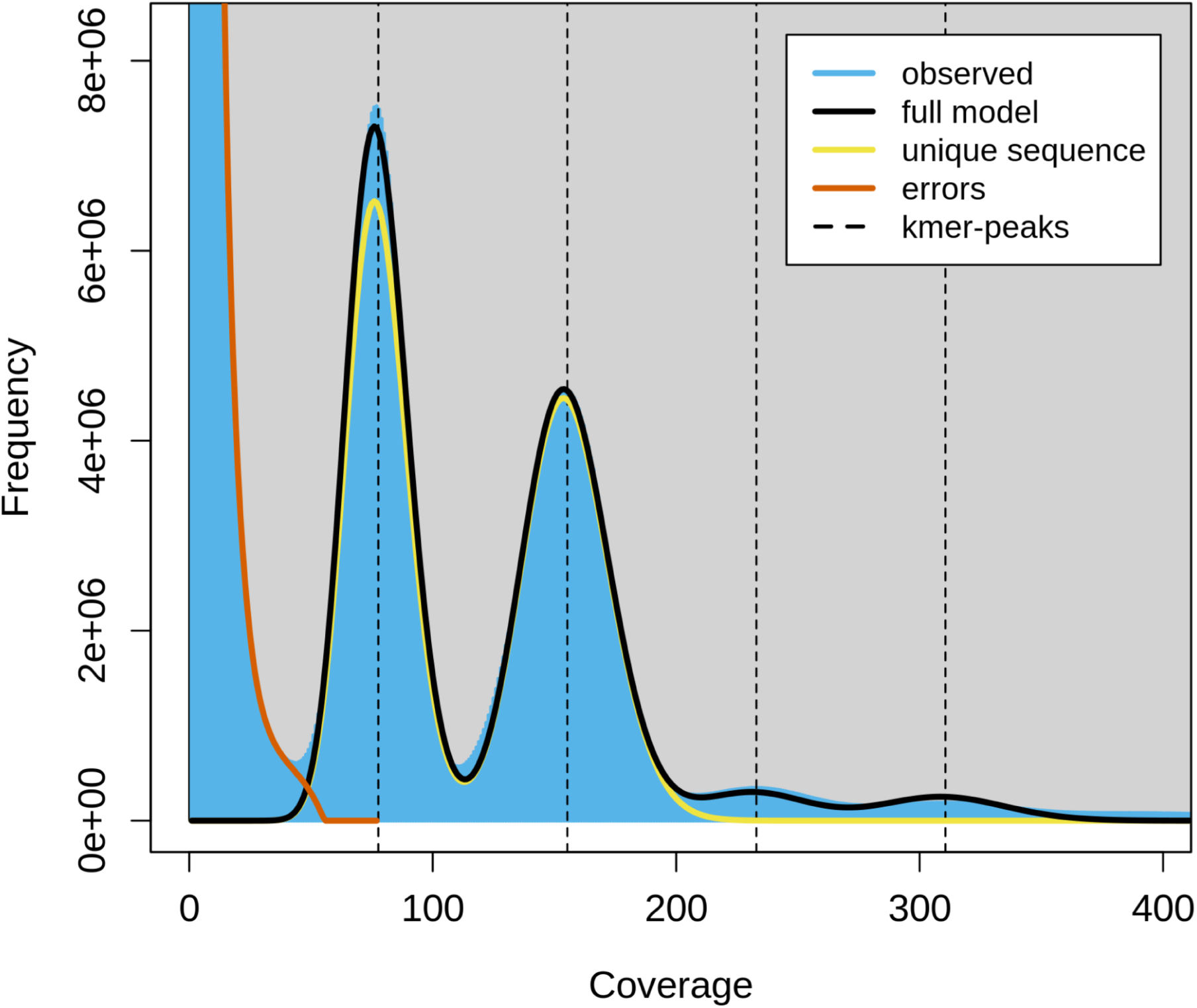
Genome size estimation plot of the *Porites harrisoni* genome. Genome size estimation plot based on k-mer distribution (k = 21) using Meryl v1.3 and GenomeScope 2.0, based on ONT reads with a low error rate (1.02%). The analysis revealed a diploid genome (p = 2) with a size of 599 Mb that is largely unique (50.3%) and predominantly homozygous (98% AA), with very little heterozygosity (1.96% AB).

The initial draft assembly with very high completeness (see ‘Data validation’) was decontaminated (i.e., contigs mapping to non-cnidarian species were removed; Fig. 3), polished, and contigs shorter than 200 bp were removed to improve accuracy and eliminate spurious fragments. Additionally, the mitogenome of *P. harrisoni* was assembled as a single-contig using Canu [57] and circularized using Circlator v1.5.5 [58], resulting in a total length of 18,639 bp closely matching the published *P. harrisoni* mitogenome from the southern Red Sea [54].

**Figure 3.**
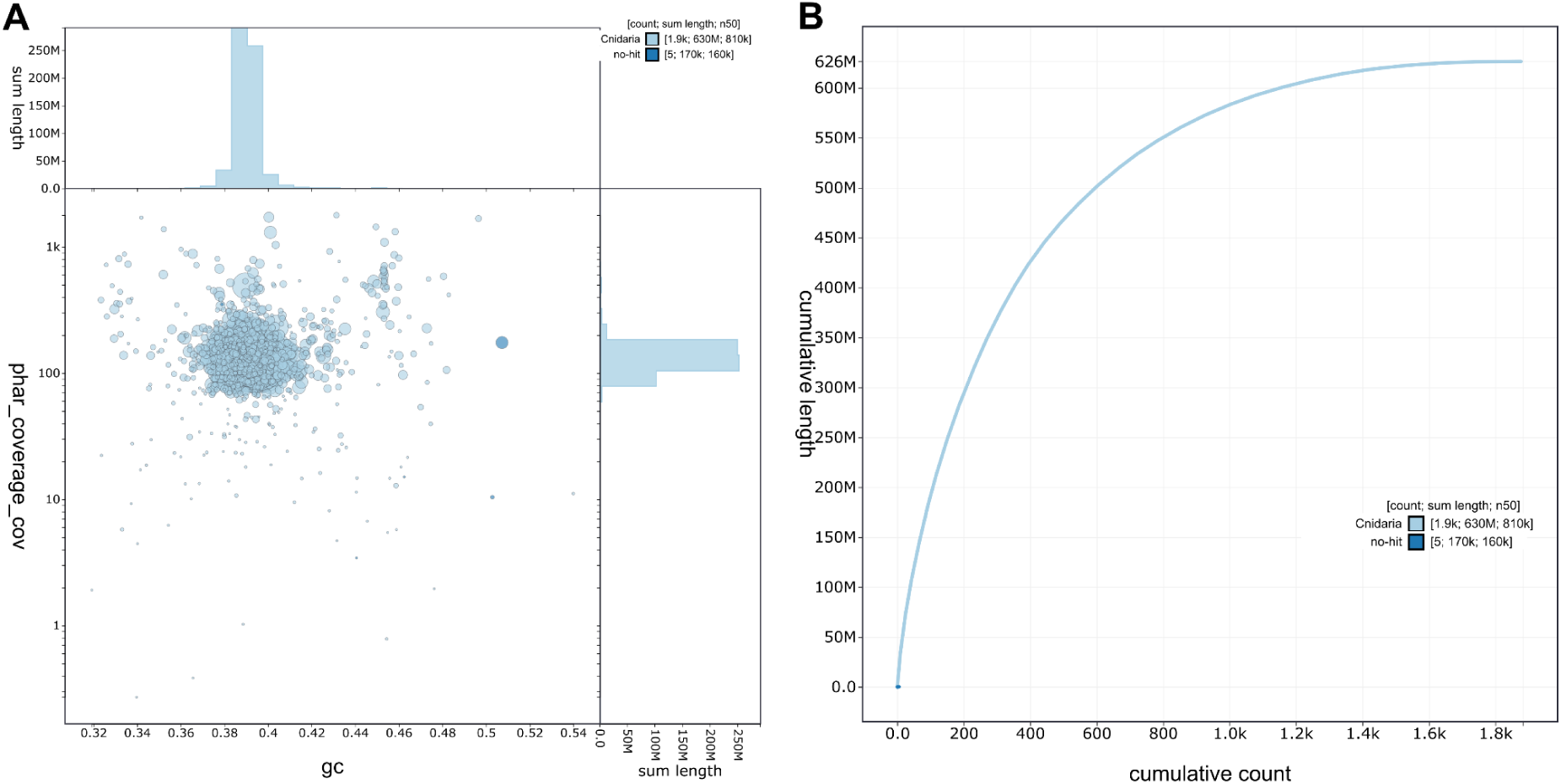
Quality assessment of the *Porites harrisoni* genome assembly using BlobToolKit v4.3.0. **(A)** GC-coverage plot showing sequence coverage (vertical axis) and GC content (horizontal axis) following decontamination (i.e., removal of contigs mapping to non-cnidarian species). Contigs are colored by phylum (light blue = Cnidaria, dark blue = no hit); circles are sized in proportion to contig length. Histograms show the distribution of contig length sums along each axis. **(B)** Cumulative sequence length of the final assembly after removal of non-target species contigs. Curves show cumulative length for all contigs with colors denoting phylum (light blue = Cnidaria, dark blue = no hit).

Lastly, the final genome assembly (nuclear genome and mitogenome) covered 1,883 contigs with a total length of 626.7 Mb and a contig N50 of 807.4 kb (Fig. 4; Table 2). Repetitive sequences identified through a combination of RepeatModeler and EDTA [61, 62] comprised a total of 371,184,681 bp (59.23% of the assembled genome), including retroelements (15.89%), DNA transposons (10.00%), and unclassified repeats (31.71%; Table 3). This is higher than for other *Porites* genomes, which have between 40 to 50% repeat coverage [35], and is likely due to the combined use of multiple repeat-identification methods rather than a single approach.

**Figure 4.**
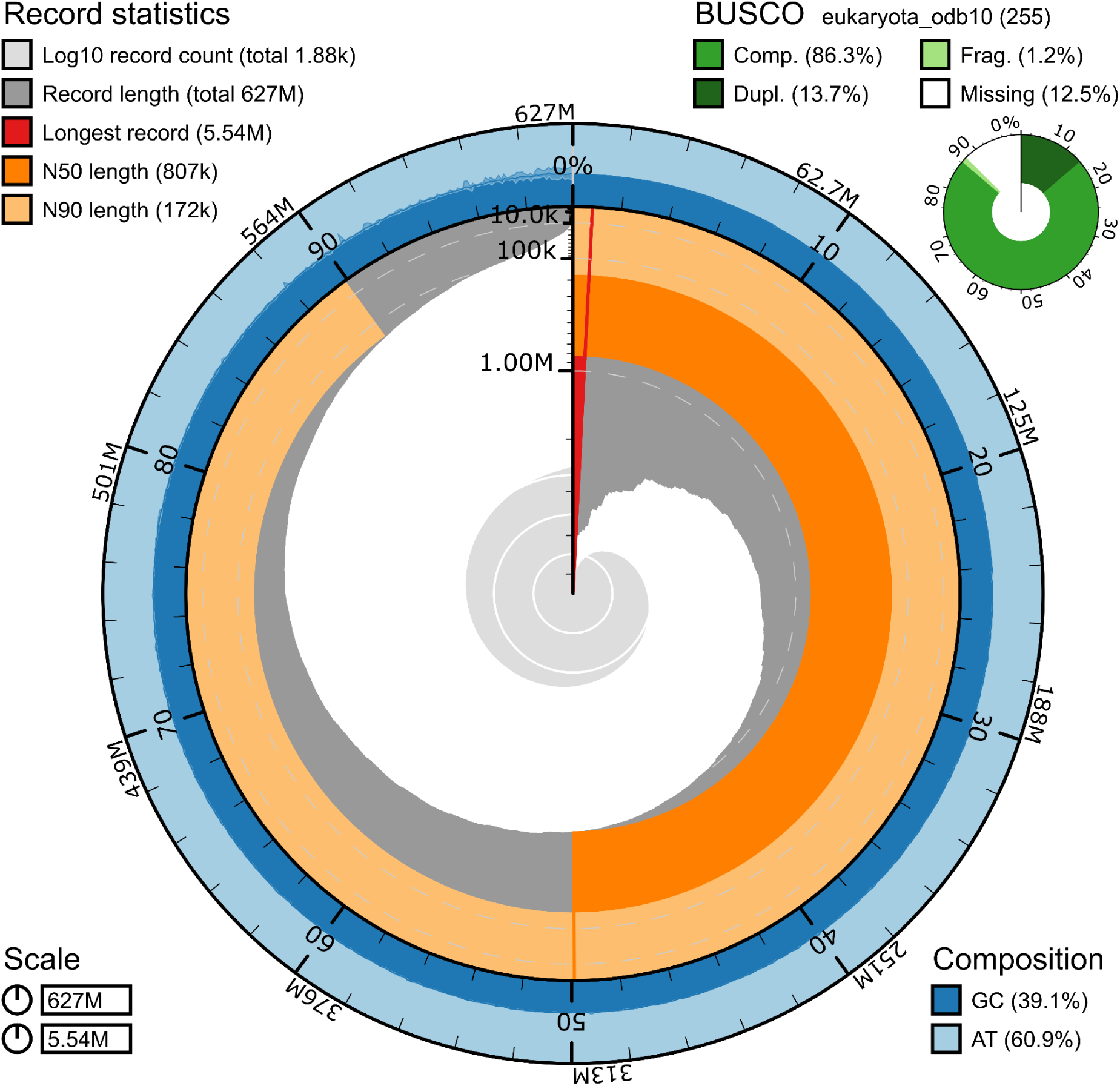
Assembly of the *Porites harrisoni* genome. The snail plot generated by BlobToolKit v4.3.0 is divided into 1,000 size-ordered bins around the circumference, with each bin representing 0.1% of the 626,672,702 bp assembly. The distribution of sequence lengths is shown in dark grey with the plot radius scaled to the longest sequence present in the assembly (5,540,956 bp, shown in red). Orange and pale-orange arcs show the N50 (807,480 bp) and N90 (172,108 bp) sequence lengths, respectively. The pale grey spiral shows the cumulative sequence count on a log scale, with white scale lines showing successive orders of magnitude. The blue and pale-blue area around the outside of the plot shows the distribution of GC, AT, and N percentages in the same bins as the inner plot. A summary of complete, fragmented, duplicated, and missing BUSCO genes in the eukaryota_odb10 set is shown at the top right (Table 2).

**Table 2.**
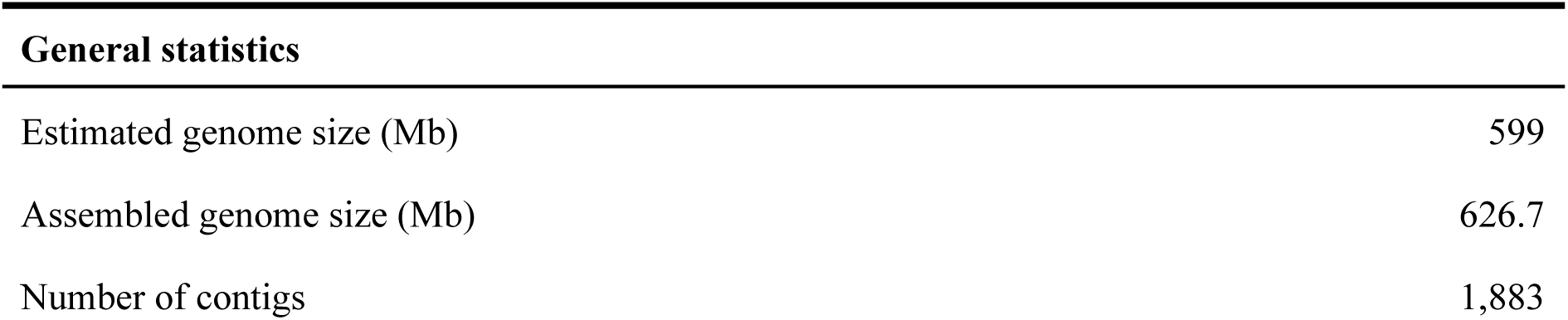
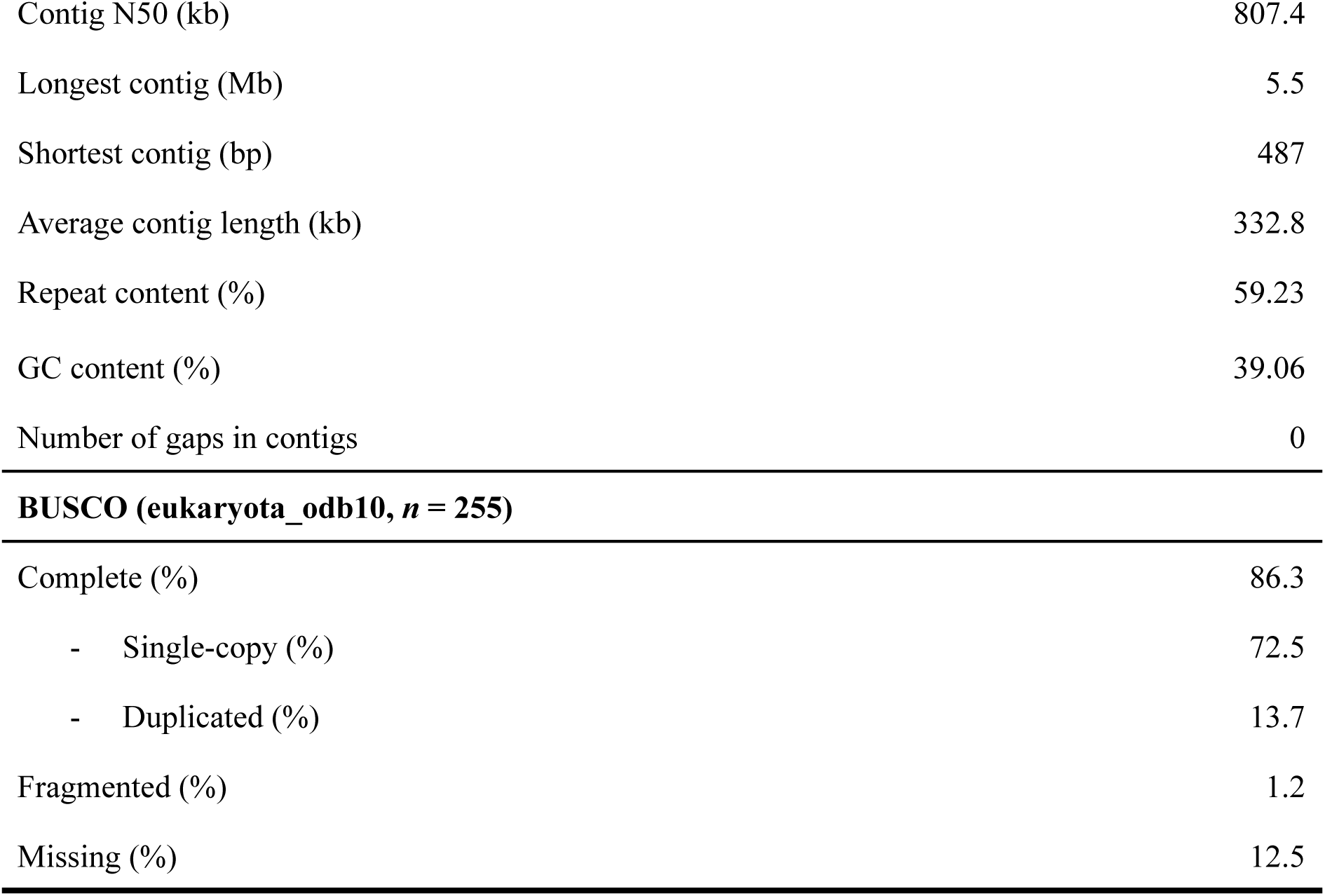
Genome assembly statistics of *Porites harrisoni*.

| General statistics |  |
| --- | --- |
| Estimated genome size (Mb) | 599 |
| Assembled genome size (Mb) | 626.7 |
| Number of contigs | 1,883 |
| Contig N50 (kb) | 807.4 |
| Longest contig (Mb) | 5.5 |
| Shortest contig (bp) | 487 |
| Average contig length (kb) | 332.8 |
| Repeat content (%) | 59.23 |
| GC content (%) | 39.06 |
| Number of gaps in contigs | 0 |
| **BUSCO (eukaryota odb10, $n = 255$)** |  |
| Complete (%) | 86.3 |
| - Single-copy (%) | 72.5 |
| - Duplicated (%) | 13.7 |
| Fragmented (%) | 1.2 |
| Missing (%) | 12.5 |

**Table 3.**
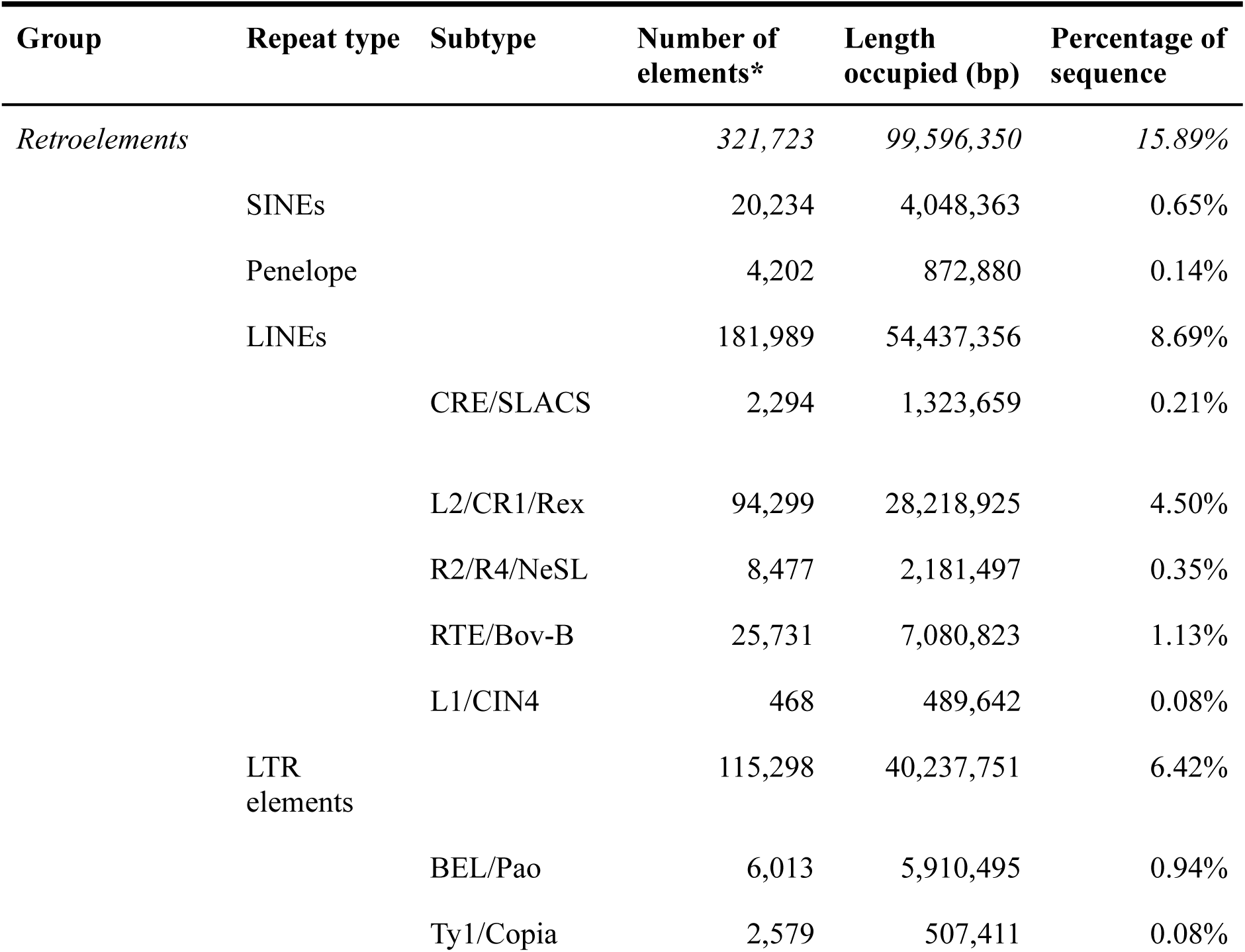
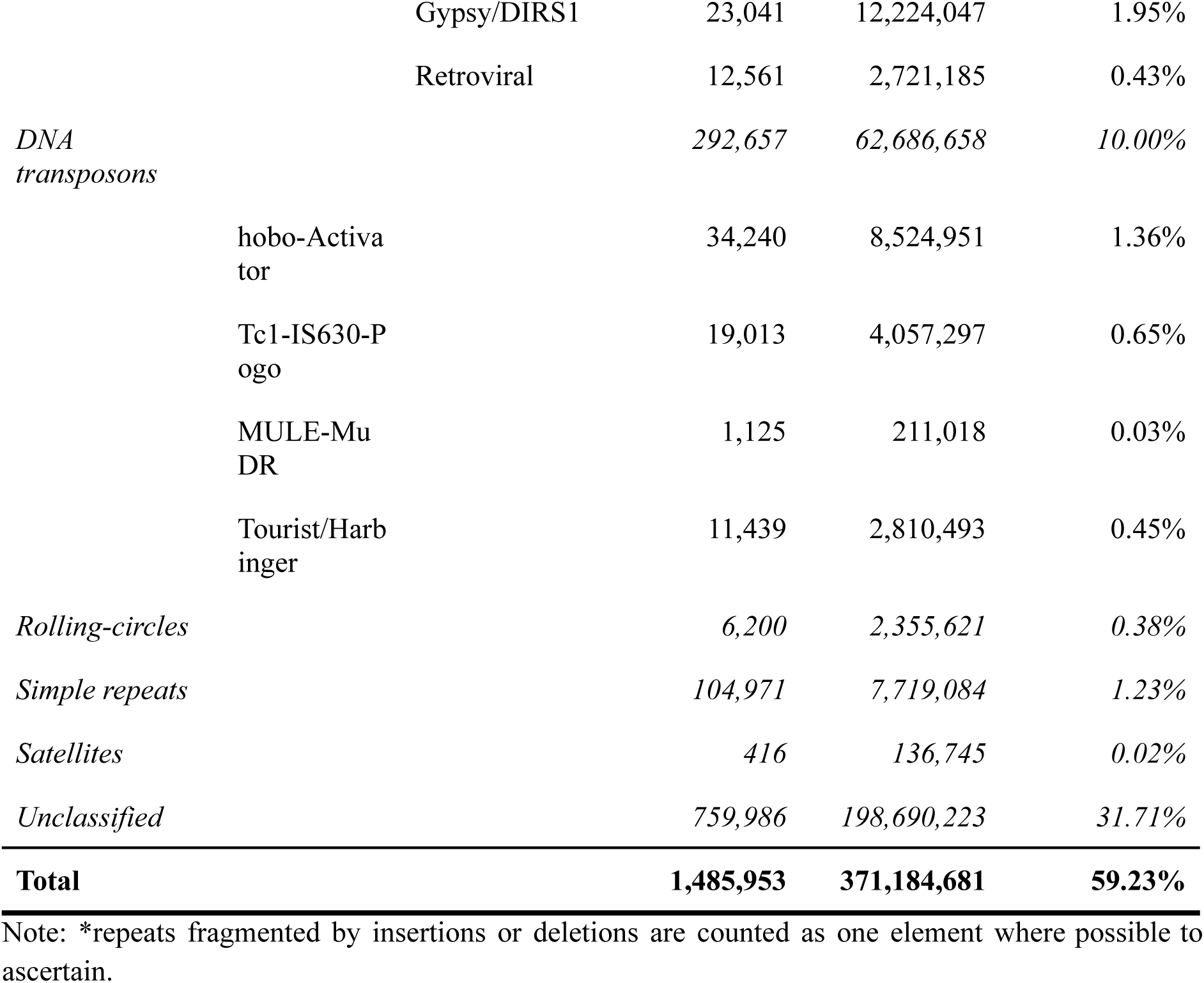
Repetitive content of the *Porites harrisoni* genome assembly.

### Genome annotation

The *Porites harrisoni* genome assembly was annotated using the BRAKER3 pipeline [64, 88] and functional annotation was done using funannotate [52]. The resulting annotation included 27,823 protein-coding genes and 674 high-confidence eukaryotic tRNAs. The average gene length was 10,193 bp, and the average protein length was 479 amino acids (aa; Table 4). The number of protein-coding genes is lower than in other *Porites* genomes (between 30,000 to 40,000 genes; [13, 35]), but is comparable to other coral genomes from the complex and robust clade [32, 89–91]. BUSCO completeness comparisons indicate a higher proportion of missing conserved genes in the *P. harrisoni* assembly (12.5% versus ∼1–3% in other published *Porites* genomes), suggesting that some genes may be absent due to assembly fragmentation, limited sequencing depth, or sequencing technology differences (e.g., short-read, long-read, hybrid assembly).

**Table 4.**
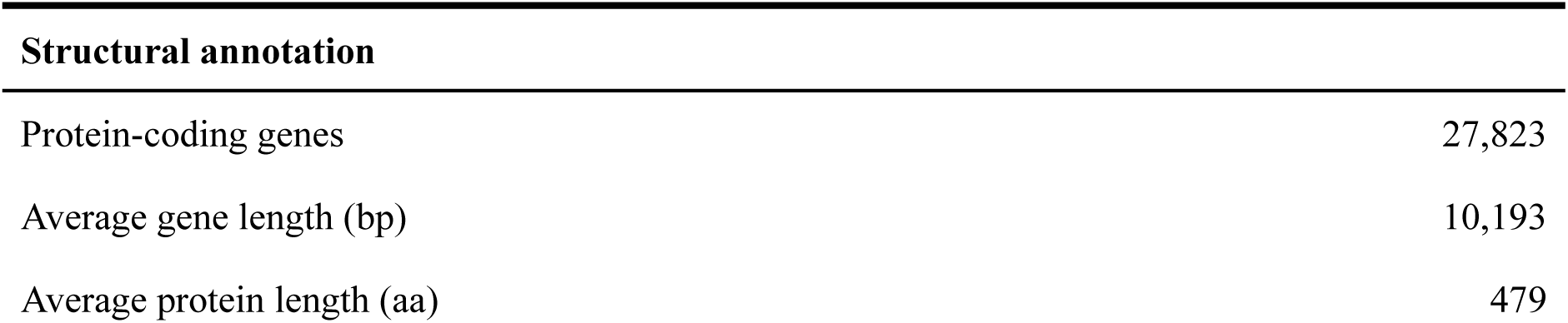
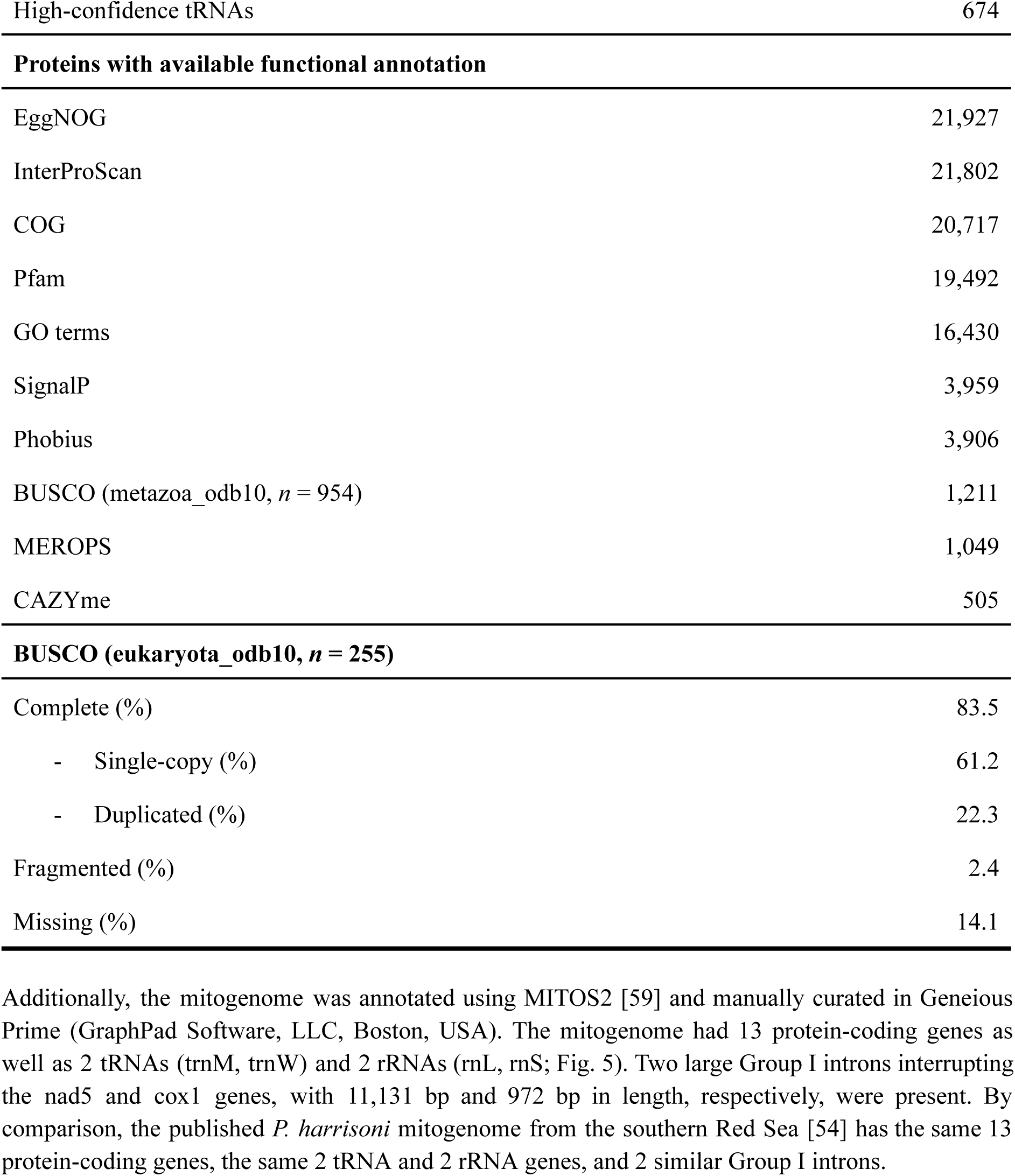
Genome annotation statistics for *Porites harrisoni*.

Additionally, the mitogenome was annotated using MITOS2 [59] and manually curated in Geneious Prime (GraphPad Software, LLC, Boston, USA). The mitogenome had 13 protein-coding genes as well as 2 tRNAs (trnM, trnW) and 2 rRNAs (rnL, rnS; Fig. 5). Two large Group I introns interrupting the nad5 and cox1 genes, with 11,131 bp and 972 bp in length, respectively, were present. By comparison, the published *P. harrisoni* mitogenome from the southern Red Sea [54] has the same 13 protein-coding genes, the same 2 tRNA and 2 rRNA genes, and 2 similar Group I introns.

**Figure 5:**
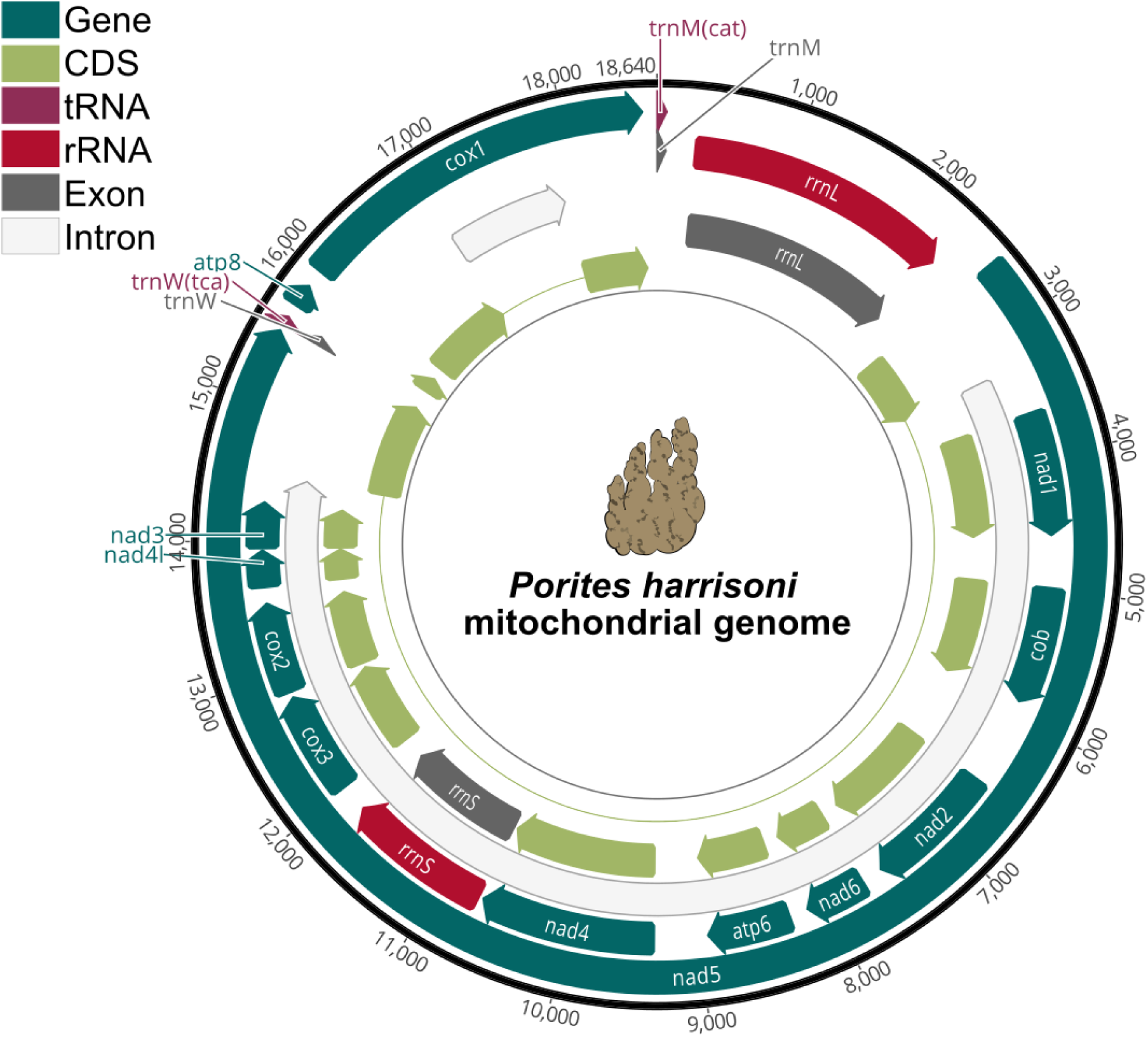
Mitochondrial genome (mitogenome) of *Porites harrisoni*. The read orientation of the respective genes is indicated by the direction of the arrows. The annotated features are demarcated by color: teal indicates the annotated genes (*n* = 13), light green denotes their corresponding coding sequences (CDS); tRNAs (*n* = 2) are in dark pink, rRNAs (*n* = 2) in red, and their respective exons are in grey (*n* = 4); introns interrupting the nad5 and cox1 genes (*n* = 2) are in white.

### Data validation

The initial draft assembly of the *Porites harrisoni* genome was 792 Mb and exhibited a high level of completeness, with a BUSCO score of 98.0% (single = 81.1%, duplicated = 16.9%, fragmented = 1.2%, missing = 0.8%) based on the eukaryota_odb10 reference set (*n* = 255), indicating that the vast majority of expected single-copy orthologs were successfully recovered. Following the removal of contigs not aligning to cnidarian sequences to ensure taxonomic accuracy and removal of non-coral sequences, the completeness was reassessed using BUSCO, resulting in a completeness score of 86.3% (single = 72.5%, duplicated = 13.7%, fragmented = 1.2%, missing = 12.5%) utilizing the eukaryota_odb10 reference set (*n* = 255; Table 2). Further refinements of the *P. harrisoni* assembly could benefit from incorporating additional long-read (e.g., PacBio HiFi) and/or short-read resequencing data to improve contiguity and resolution, especially with regard to unequivocally resolving the definitive number of protein-coding genes. The integration of Hi-C data could aid in the generation of a chromosome-level assembly (e.g., [92]), albeit misjoints could be introduced if Hi-C data is not of sufficient quality. Subsequent evaluation of completeness based on the set of annotated protein-coding genes yielded a BUSCO completeness of 83.5% (single = 61.2%, duplicated = 22.3%, fragmented = 2.4%, missing = 14.1%) using the eukaryota_odb10 reference set (*n* = 255; Table 4). Although other *Porites* genomes show higher completeness scores [13, 35], the results in this study confirm that the final *Porites harrisoni* assembly and annotation represent a high-quality and taxonomically validated coral genome, with the reduction in BUSCO completeness attributable to the deliberate exclusion of non-cnidarian sequences, rather than a reduction in genomic accuracy.

## Conclusion

The genome presented in this study represents the first assembled genome of *Porites harrisoni* from the thermally extreme environment of the southern PAG. As one of the world’s most thermally resilient coral species, the genome of this coral provides a critical reference for investigating the genomic basis of heat tolerance, rapid adaptation, and host-algal symbiosis under severe thermal stress. The availability of a high-quality and complete mitochondrial genome from the PAG enables comparative genomic analyses among coral species and populations, facilitating the identification of adaptive loci. Placed alongside the increasing suite of high-quality reference genomes for corals, e.g., [32, 35, 89–91, 93] and other non-model taxa [94–96], this genome strengthens the expanding genomic foundation that is driving advances in evolutionary biology and conservation science. Moreover, this genome serves as a foundation for future studies on coral evolutionary genomics and population structure of *P. harrisoni*, and will support studies aimed at predicting coral responses to ongoing ocean warming.

## Availability of Source Code

All bioinformatic tools used in this study can be found in Table 5; all curated pipelines are available on GitHub under https://github.com/SequAna-Ukon/Porites_genome.

**Table 5.**
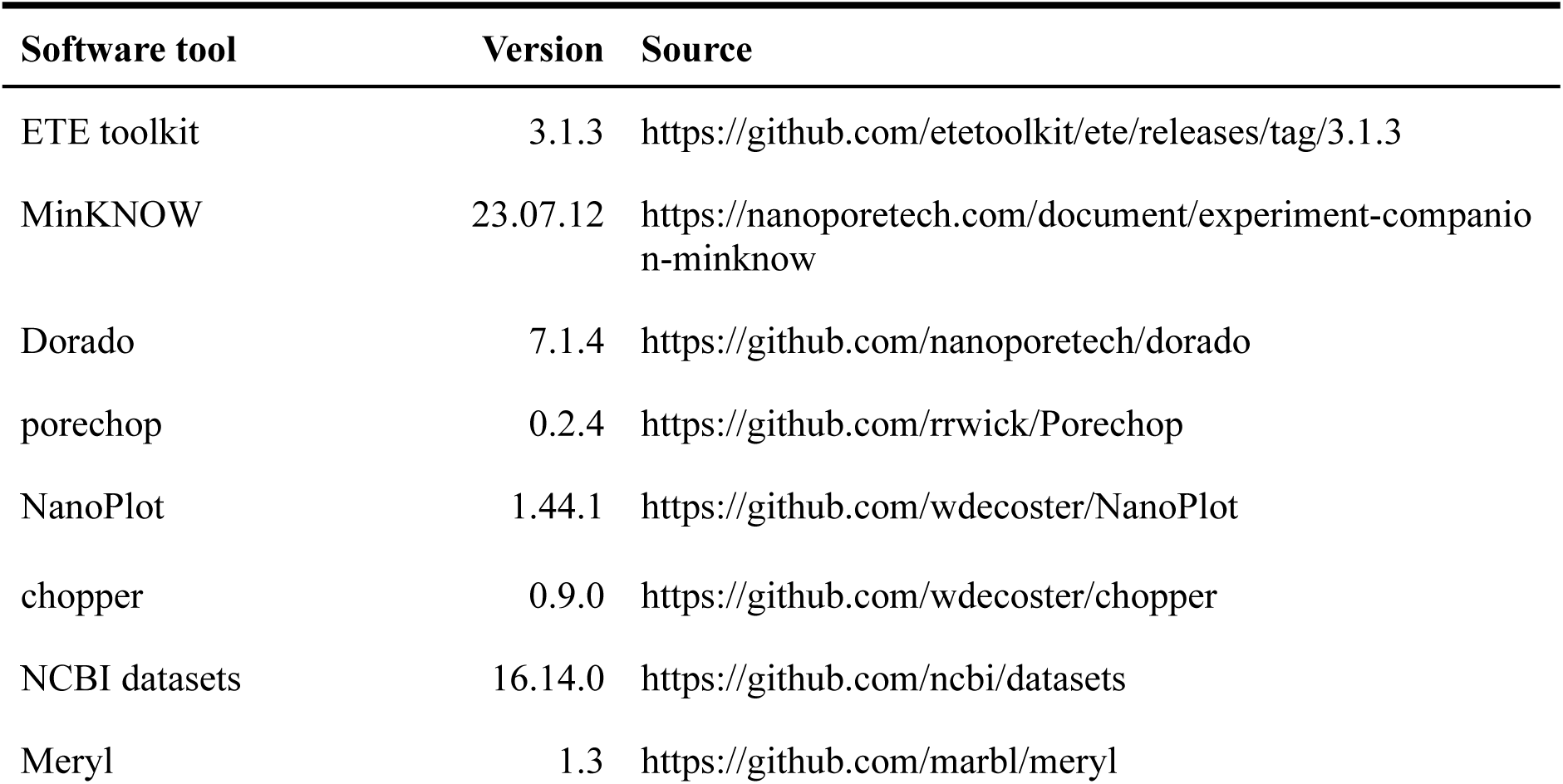
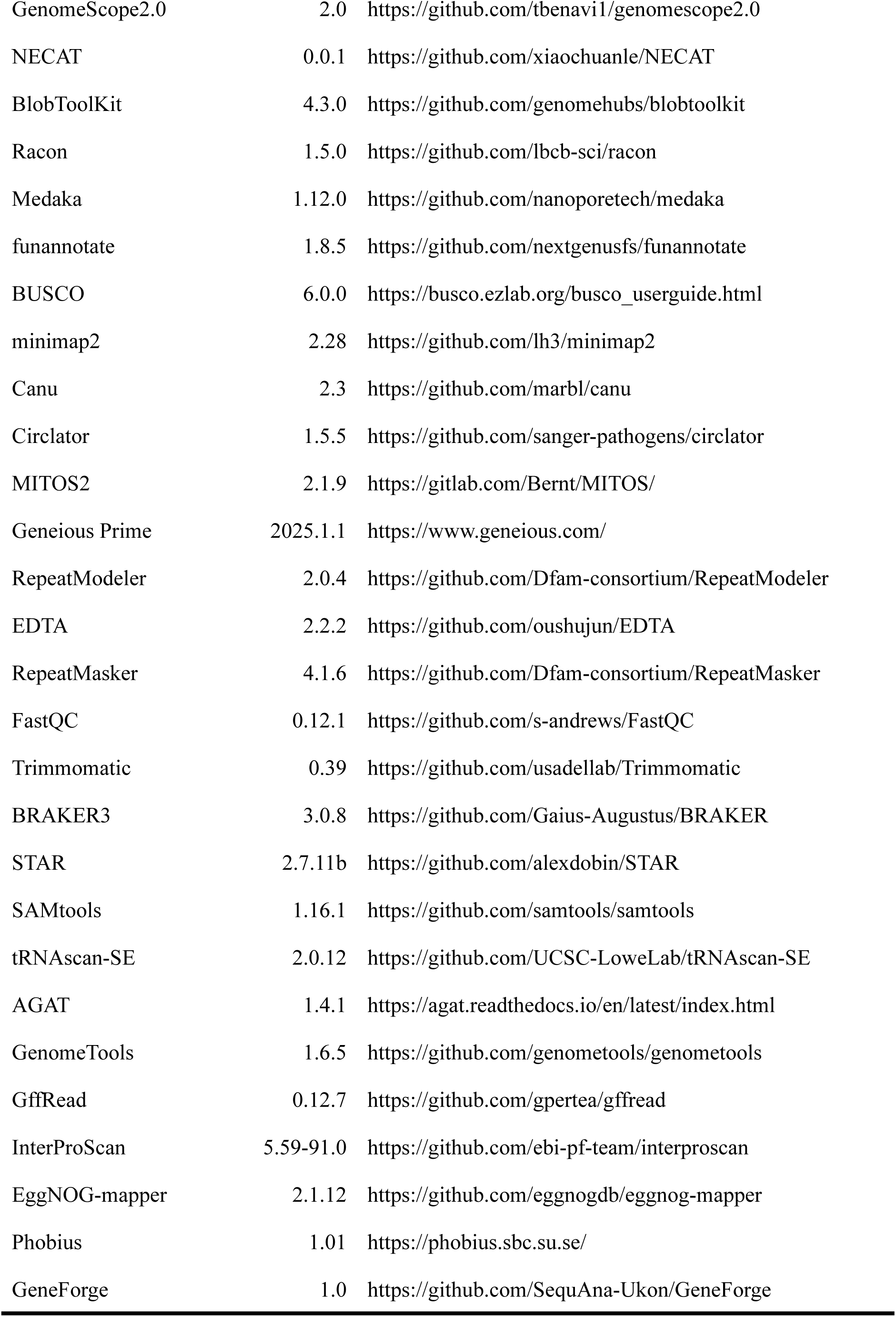
Software tools: versions and sources.

Project name: Porites_genome

Project home page: https://github.com/SequAna-Ukon/Porites_genome

Operating system: Platform dependent

Programming language: bash

License: MIT

## Data Availability

Raw gDNA sequencing data are deposited at NCBI under the BioProject PRJNA1111311 (https://www.ncbi.nlm.nih.gov/bioproject/PRJNA1111311) and raw RNA-Seq data are deposited under the BioProject PRJNA1354406 (https://www.ncbi.nlm.nih.gov/bioproject/PRJNA1354406), both accessible under the Umbrella BioProject PRJNA749006 (https://www.ncbi.nlm.nih.gov/bioproject/PRJNA749006). This Whole Genome Shotgun project has been deposited at DDBJ/ENA/GenBank under the accession JBDLLT000000000 (https://www.ncbi.nlm.nih.gov/nuccore/JBDLLT000000000). The version described in this paper is version JBDLLT020000000. Assembly and annotation files can be downloaded via https://www.ncbi.nlm.nih.gov/datasets/genome/GCA_040938025.2/.

## Declarations

### List of abbreviations

bp: base pair
BUSCO: benchmarking universal single-copy orthologs
CBASS: Coral Bleaching Automated Stress System
CDS: coding sequence
GO: Gulf of Oman
kb: kilobase pairs
LINEs: long interspersed nuclear elements
LTRs: long terminal repeats
Mb: megabase pairs
NCBI: National Center for Biotechnology Information
ONT: Oxford Nanopore Technologies
PAG: Persian/Arabian Gulf
SINEs: short interspersed nuclear elements
UAE: United Arab Emirates
UTR: untranslated region

### Ethics approval & consent for publication

Not applicable.

## Acknowledgements

We thank the NYU Abu Dhabi Core Technology Marine Science group and particularly Dain McParland for field support, and we thank Luigi Colin for technical support. We gratefully acknowledge the Environment Agency Abu Dhabi research permit with the reference number 56930. We thank the NGS Competence Center Tübingen (NCCT) for their sequencing services and technical assistance (project number QCVTS-SEQ1077).

## Author contributions

CRV conceived and conceptualized the study. AF, GP, HM, RA, and CRV collected the samples. JAB provided field support. AF and RA processed the samples. AF and AS assembled and annotated the genome. AF visualized the results. AF, AS, and CRV interpreted all analyses and results. AF, AS, RA, and CRV wrote the manuscript draft. All authors edited and approved the final manuscript.

## References

1. Riegl, B. M., & Purkis, S. J. (2012). Coral reefs of the gulf: Adaptation to climatic extremes in the world’s hottest sea. In Coral Reefs of the World (pp. 1–4). Dordrecht: Springer Netherlands. 10.1007/978-94-007-3008-3_1

2. Burt, J. A., Camp, E. F., Enochs, I. C., Johansen, J. L., Morgan, K. M., Riegl, B., & Hoey, A. S. (2020). Insights from extreme coral reefs in a changing world. Coral reefs, 39(3), 495–507. 10.1007/s00338-020-01966-y

3. Camp, E. F., Suggett, D. J., Pogoreutz, C., Nitschke, M. R., Houlbreque, F., Hume, B. C. C. et al. (2020). Corals exhibit distinct patterns of microbial reorganisation to thrive in an extreme inshore environment. Coral Reefs, 39(3), 701–716. 10.1007/s00338-019-01889-3

4. Coles, S. L., & Riegl, B. M. (2013). Thermal tolerances of reef corals in the Gulf: a review of the potential for increasing coral survival and adaptation to climate change through assisted translocation. Marine Pollution Bulletin, 72(2), 323–332. 10.1016/j.marpolbul.2012.09.006

5. Vaughan, G. O., Al-Mansoori, N., & Burt, J. A. (2018). Chapter 1 - The Arabian Gulf (pp. 1–23). Elsevier Ltd. 10.1016/B978-0-08-100853-9.00001-4

6. Hume, B. C. C., D’Angelo, C., Smith, E. G., Stevens, J. R., Burt, J., & Wiedenmann, J. (2015). Symbiodinium thermophilum sp. nov., a thermotolerant symbiotic alga prevalent in corals of the world’s hottest sea, the Persian/Arabian Gulf. Scientific Reports, 5(1), 8562. 10.1038/srep08562

7. Burt, J. A. (2024). Coral reefs of the Emirates. In A Natural History of the Emirates (pp. 325–351). Cham: Springer Nature Switzerland. 10.1007/978-3-031-37397-8_11

8. Bhattacharya, D., Agrawal, S., Aranda, M., Baumgarten, S., Belcaid, M., Drake, J. L. et al. (2016). Comparative genomics explains the evolutionary success of reef-forming corals. eLife, 5, e13288. 10.7554/eLife.13288

9. Romano, S. L., & Palumbi, S. R. (1996). Evolution of Scleractinian Corals Inferred from Molecular Systematics. Science, 271(5249), 640–642. 10.1126/science.271.5249.640

10. Baird, A. H., & Marshall, P. A. (2002). Mortality, growth and reproduction in scleractinian corals following bleaching on the Great Barrier Reef. Marine Ecology Progress Series, 237, 133–141. 10.3354/meps237133

11. Coles, S. L., & Brown, B. E. (2003). Coral bleaching—capacity for acclimatization and adaptation. Advances in Marine Biology. 10.1016/S0065-2881(03)46004-5

12. Burt, J. A., Paparella, F., Al-Mansoori, N., Al-Mansoori, A., & Al-Jailani, H. (2019). Causes and consequences of the 2017 coral bleaching event in the southern Persian/Arabian Gulf. Coral Reefs, 38(4), 567–589. 10.1007/s00338-019-01767-y

13. Shinzato, C., Takeuchi, T., Yoshioka, Y., Tada, I., Kanda, M., Broussard, C., … Inoue, M. (2021). Whole-genome sequencing highlights conservative genomic strategies of a stress-tolerant, long-lived scleractinian coral, Porites australiensis Vaughan, 1918. Genome Biology and Evolution, 13(12), evab270. 10.1093/gbe/evab270

14. Alderdice, R., Hume, B. C. C., Kühl, M., Pernice, M., Suggett, D. J., & Voolstra, C. R. (2022). Disparate inventories of hypoxia gene sets across corals align with inferred environmental resilience. Frontiers in Marine Science, 9. 10.3389/fmars.2022.834332

15. Riegl, B. M., Benzoni, F., Samimi-Namin, K., & Sheppard, C. (2012). The hermatypic scleractinian (hard) coral fauna of the gulf. In Coral Reefs 2of the World (pp. 187–224). Dordrecht: Springer Netherlands. 10.1007/978-94-007-3008-3_11

16. Smith, E. G., Hume, B. C. C., Delaney, P., Wiedenmann, J., & Burt, J. A. (2017). Genetic structure of coral-Symbiodinium symbioses on the world’s warmest reefs. PloS One, 12(6), e0180169. 10.1371/journal.pone.0180169

17. Howells, E. J., Bauman, A. G., Vaughan, G. O., Hume, B. C. C., Voolstra, C. R., & Burt, J. A. (2020). Corals in the hottest reefs in the world exhibit symbiont fidelity not flexibility. Molecular Ecology, 29(5), 899–911. 10.1111/mec.15372

18. Hume, B. C. C., Voolstra, C. R., Arif, C., D’Angelo, C., Burt, J. A., Eyal, G., … Wiedenmann, J. (2016). Ancestral genetic diversity associated with the rapid spread of stress-tolerant coral symbionts in response to Holocene climate change. Proceedings of the National Academy of Sciences of the United States of America, 113(16), 4416–4421. 10.1073/pnas.1601910113

19. Smith, E. G., Hazzouri, K. M., Choi, J. Y., Delaney, P., Al-Kharafi, M., Howells, E. J. et al. (2022). Signatures of selection underpinning rapid coral adaptation to the world’s warmest reefs. Science Advances, 8(2), eabl7287. 10.1126/sciadv.abl7287

20. Howells, E. J., Abrego, D., Liew, Y. J., Burt, J. A., Meyer, E., & Aranda, M. (2021). Enhancing the heat tolerance of reef-building corals to future warming. Science Advances, 7(34). 10.1126/sciadv.abg6070

21. Howells, E. J., Abrego, D., Schmidt-Roach, S., Puill-Stephan, E., Denis, H., Harii, S. et al. (2025). Marine heatwaves select for thermal tolerance in a reef-building coral. Nature Climate Change, 15(8), 829–832. 10.1038/s41558-025-02381-3

22. Riegl, B. M., Purkis, S. J., Al-Cibahy, A. S., Al-Harthi, S., Grandcourt, E., Al-Sulaiti, K. et al. (2012). Coral bleaching and mortality thresholds in the SE gulf: Highest in the world. In Coral Reefs of the World (pp. 95–105). Dordrecht: Springer Netherlands. 10.1007/978-94-007-3008-3_6

23. Kinsman, D. J. J. (1964). Reef coral tolerance of high temperatures and salinities. Nature, 202(4939), 1280–1282. 10.1038/2021280a0

24. Sheppard, C. R. C. (1988). Similar trends, different causes: Responses of corals to stressed environments in Arabian Seas. Proceedings of the 6th International Coral Reef Symposium Townsville, Australia, 3, 297–302.

25. Alderdice, R., Perna, G., Manns, H., Fiesinger, A., Colin, L., Stankiewicz, K. et al. (2025). Summer temperature maxima already challenge thermal capacities of coral in the Southern Persian/Arabian Gulf. Research Square. 10.21203/rs.3.rs-8231598/v1

26. McCartin, L., Saso, E., Vohsen, S. A., Pittoors, N., Demetriades, P., McFadden, C. S. et al. (2024). Nuclear eDNA metabarcoding primers for anthozoan coral biodiversity assessment. PeerJ, 12, e18607. 10.7717/peerj.18607

27. Banguera-Hinestroza, E., Sawall, Y., Al-Sofyani, A., Mardulyn, P., Fuertes-Aguilar, J., Cárdenas-Henao, H., … Flot, J.-F. (2018). mtDNA recombination indicative of hybridization suggests a role of the mitogenome in the adaptation of reef-building corals to extreme environments. Evolutionary Biology. bioRxiv. Retrieved from https://www.biorxiv.org/content/10.1101/462069v3

28. Putnam, N. H., Srivastava, M., Hellsten, U., Dirks, B., Chapman, J., Salamov, A. et al. (2007). Sea anemone genome reveals ancestral eumetazoan gene repertoire and genomic organization. Science (New York, N.Y.), 317(5834), 86–94. 10.1126/science.1139158

29. Baumgarten, S., Simakov, O., Esherick, L. Y., Liew, Y. J., Lehnert, E. M., Michell, C. T. et al. (2015). The genome of Aiptasia, a sea anemone model for coral symbiosis. Proceedings of the National Academy of Sciences of the United States of America, 112(38), 11893–11898. 10.1073/pnas.1513318112

30. Liew, Y. J., Howells, E. J., Wang, X., Michell, C. T., Burt, J. A., Idaghdour, Y., & Aranda, M. (2020). Intergenerational epigenetic inheritance in reef-building corals. Nature Climate Change, 10(3), 254–259. 10.1038/s41558-019-0687-2

31. Cunning, R., Bay, R. A., Gillette, P., Baker, A. C., & Traylor-Knowles, N. (2018). Comparative analysis of the Pocillopora damicornis genome highlights role of immune system in coral evolution. Scientific Reports, 8(1), 16134. 10.1038/s41598-018-34459-8

32. Voolstra, C. R., Li, Y., Liew, Y. J., Baumgarten, S., Zoccola, D., Flot, J.-F. et al. (2017). Comparative analysis of the genomes of Stylophora pistillata and Acropora digitifera provides evidence for extensive differences between species of corals. Scientific Reports, 7(1), 17583. 10.1038/s41598-017-17484-x

33. Fuller, Z. L., Mocellin, V. J. L., Morris, L. A., Cantin, N., Shepherd, J., Sarre, L. et al. (2020). Population genetics of the coral Acropora millepora: Toward genomic prediction of bleaching. Science (New York, N.Y.), 369(6501), eaba4674. 10.1126/science.aba4674

34. Helmkampf, M., Bellinger, M. R., Geib, S. M., Sim, S. B., & Takabayashi, M. (2019). Draft genome of the rice coral Montipora capitata obtained from linked-read sequencing. Genome Biology and Evolution, 11(7), 2045–2054. 10.1093/gbe/evz135

35. Noel, B., Denoeud, F., Rouan, A., Buitrago-López, C., Capasso, L., Poulain, J. et al. (2023). Pervasive tandem duplications and convergent evolution shape coral genomes. Genome Biology, 24(1). 10.1186/s13059-023-02960-7

36. Huerta-Cepas, J., Serra, F., & Bork, P. (2016). ETE 3: Reconstruction, analysis, and visualization of phylogenomic data. Molecular Biology and Evolution, 33(6), 1635–1638. 10.1093/molbev/msw046

37. Romano, S. L., & Cairns, S. (2000). Molecular phylogenetic hypotheses for the evolution of scleractinian corals. Bulletin of Marine Science, 67(3), 1043–1068.

38. Vaga, C. F., Quattrini, A. M., Galvão de Lossio E Seiblitz, I., Huang, D., Quek, Z. B. R., Stolarski, J. et al. (2025). A global coral phylogeny reveals resilience and vulnerability through deep time. Nature, 648(8093), 377–382. 10.1038/s41586-025-09615-6

39. Evensen, N. R., Parker, K. E., Oliver, T. A., Palumbi, S. R., Logan, C. A., Ryan, J. S. et al. (2023). The Coral Bleaching Automated Stress System (CBASS): A low-cost, portable system for standardized empirical assessments of coral thermal limits. *Limnology and Oceanography*, Methods, 21(7), 421–434. 10.1002/lom3.10555

40. Baums, I., & Kitchen, S. A. (2020). Acropra DNA extraction with Qiagen DNAease tissue kit v1. Protocolsio. 10.17504/protocols.io.bcwuixew

41. Oxford Nanopore Technologies. (2024). Dorado. Retrieved from https://github.com/nanoporetech/dorado

42. Wick, R. (2018). Porechop. Retrieved from https://github.com/rrwick/Porechop

43. De Coster, W., & Rademakers, R. (2023). NanoPack2: population-scale evaluation of long-read sequencing data. *Bioinformatics (Oxford*, England), 39(5). 10.1093/bioinformatics/btad311

44. O’Leary, N. A., Cox, E., Holmes, J. B., Anderson, W. R., Falk, R., Hem, V. et al. (2024). Exploring and retrieving sequence and metadata for species across the tree of life with NCBI Datasets. Scientific Data, 11(1), 732. 10.1038/s41597-024-03571-y

45. Voolstra, C. R., Perna, G., & Alderdice, R. (2022). RNA preservation & RNA extraction protocol suitable for field collection of coral samples. Zenodo. 10.5281/zenodo.6492452

46. Rhie, A., Walenz, B. P., Koren, S., & Phillippy, A. M. (2020). Merqury: reference-free quality, completeness, and phasing assessment for genome assemblies. Genome Biology, 21(1), 245. 10.1186/s13059-020-02134-9

47. Ranallo-Benavidez, T., Jaron, K., Jaron, K., Schatz, M., & Schatz, M. (2020). GenomeScope 2.0 and Smudgeplot for reference-free profiling of polyploid genomes. Nature Communications, 11. 10.1038/s41467-020-14998-3

48. Chen, Y., Nie, F., Xie, S.-Q., Zheng, Y.-F., Dai, Q., Bray, T. et al. (2021). Efficient assembly of nanopore reads via highly accurate and intact error correction. Nature Communications, 12(1), 60. 10.1038/s41467-020-20236-7

49. Challis, R., Richards, E., Rajan, J., Cochrane, G., & Blaxter, M. (2020). BlobToolKit - interactive quality assessment of genome assemblies. G3 (Bethesda, Md.), 10(4), 1361–1374. 10.1534/g3.119.400908

50. Sovic, I., Vaser, R., Sikic, M., & Nagarajan, N. (2021). Racon. Retrieved from https://github.com/lbcb-sci/racon

51. Oxford Nanopore Technologies. (2024). Medaka. Retrieved from https://github.com/nanoporetech/medaka

52. Palmer, J. M., & Stajich, J. (2020). Funannotate. 10.5281/zenodo.4054262

53. Simão, F. A., Waterhouse, R. M., Ioannidis, P., Kriventseva, E., & Zdobnov, E. (2015). BUSCO: assessing genome assembly and annotation completeness with single-copy orthologs. Bioinformatics, 31(19), 3210–3212. 10.1093/bioinformatics/btv351

54. Terraneo, T. I., Arrigoni, R., Benzoni, F., Forsman, Z. H., & Berumen, M. L. (2018). The complete mitochondrial genome of Porites harrisoni (Cnidaria: Scleractinia) obtained using next-generation sequencing. *Mitochondrial DNA. Part B*, Resources, 3(1), 286–287. 10.1080/23802359.2018.1443852

55. Li, H. (2017). Minimap2: pairwise alignment for nucleotide sequences. Bioinformatics, 34, 3094–3100. 10.1093/bioinformatics/bty191

56. Danecek, P., Bonfield, J. K., Liddle, J., Marshall, J., Ohan, V., Pollard, M. O. et al. (2021). Twelve years of SAMtools and BCFtools. GigaScience, 10(2), giab008. 10.1093/gigascience/giab008

57. Koren, S., Walenz, B. P., Berlin, K., Miller, J. R., Bergman, N. H., & Phillippy, A. M. (2017). Canu: scalable and accurate long-read assembly via adaptive k-mer weighting and repeat separation. Genome Research, 27(5), 722–736. 10.1101/gr.215087.116

58. Hunt, M., Silva, N. D., Otto, T. D., Parkhill, J., Keane, J. A., & Harris, S. R. (2015). Circlator: automated circularization of genome assemblies using long sequencing reads. Genome Biology, 16(1), 294. 10.1186/s13059-015-0849-0

59. Bernt, M., Donath, A., Jühling, F., Externbrink, F., Florentz, C., Fritzsch, G. et al. (2013). MITOS: improved de novo metazoan mitochondrial genome annotation. Molecular Phylogenetics and Evolution, 69(2), 313–319. 10.1016/j.ympev.2012.08.023

60. Tisthammer, K. H., Forsman, Z. H., Sindorf, V. L., Massey, T. L., Bielecki, C. R., & Toonen, R. J. (2016). The complete mitochondrial genome of the lobe coral Porites lobata (Anthozoa: Scleractinia) sequenced using ezRAD. Mitochondrial DNA. Part B, Resources, 1(1), 247–249. 10.1080/23802359.2016.1157770

61. Flynn, J. M., Hubley, R., Goubert, C., Rosen, J., Clark, A. G., Feschotte, C., & Smit, A. F. (2020). RepeatModeler2 for automated genomic discovery of transposable element families. Proceedings of the National Academy of Sciences of the United States of America, 117(17), 9451–9457. 10.1073/pnas.1921046117

62. Ou, S., Su, W., Liao, Y., Chougule, K., Agda, J. R. A., Hellinga, A. J. et al. (2019). Benchmarking transposable element annotation methods for creation of a streamlined, comprehensive pipeline. Genome Biology, 20(1), 275. 10.1186/s13059-019-1905-y

63. Nishimura, D. (2000). RepeatMasker. Biotech software & Internet report, 1(1-2), 36–39. 10.1089/152791600319259

64. Gabriel, L., Brůna, T., Hoff, K. J., Ebel, M., Lomsadze, A., Borodovsky, M., & Stanke, M. (2024). BRAKER3: Fully automated genome annotation using RNA-seq and protein evidence with GeneMark-ETP, AUGUSTUS, and TSEBRA. Genome Rresearch, 34(5), 769–777. 10.1101/gr.278090.123

65. Andrews, S. (2023). FastQC: A quality control tool for high throughput sequence data. Retrieved from https://github.com/s-andrews/FastQC

66. Bolger, A. M., Lohse, M., & Usadel, B. (2014). Trimmomatic: a flexible trimmer for Illumina sequence data. Bioinformatics, 30, 2114–2120. 10.1093/bioinformatics/btu170

67. Dobin, A., Davis, C. A., Schlesinger, F., Drenkow, J., Zaleski, C., Jha, S. et al. (2013). STAR: ultrafast universal RNA-seq aligner. *Bioinformatics (Oxford*, England), 29(1), 15–21. 10.1093/bioinformatics/bts635

68. Brůna, T., Gabriel, L., & Hoff, K. J. (2024). Navigating eukaryotic genome annotation pipelines: A route map to BRAKER, Galba, and TSEBRA. *arXiv.* Retrieved from http://arxiv.org/abs/2403.19416

69. Bruna, T., Lomsadze, A., & Borodovsky, M. (2024). A new gene finding tool GeneMark-ETP significantly improves the accuracy of automatic annotation of large eukaryotic genomes. bioRxiv. 10.1101/2023.01.13.524024

70. Buchfink, B., Xie, C., & Huson, D. H. (2015). Fast and sensitive protein alignment using DIAMOND. Nature Methods, 12(1), 59–60. 10.1038/nmeth.3176

71. Li, H. (2023). Protein-to-genome alignment with miniprot. *Bioinformatics (Oxford*, England), 39(1), btad014. 10.1093/bioinformatics/btad014

72. Huang, N., & Li, H. (2023). compleasm: a faster and more accurate reimplementation of BUSCO. *Bioinformatics (Oxford*, England), 39(10), btad595. 10.1093/bioinformatics/btad595

73. Gotoh, O. (2008). A space-efficient and accurate method for mapping and aligning cDNA sequences onto genomic sequence. Nucleic Acids Research, 36(8), 2630–2638. 10.1093/nar/gkn105

74. Iwata, H., & Gotoh, O. (2012). Benchmarking spliced alignment programs including Spaln2, an extended version of Spaln that incorporates additional species-specific features. Nucleic acids Research, 40(20), e161. 10.1093/nar/gks708

75. Kovaka, S., Zimin, A. V., Pertea, G. M., Razaghi, R., Salzberg, S. L., & Pertea, M. (2019). Transcriptome assembly from long-read RNA-seq alignments with StringTie2. Genome Biology, 20(1), 278. 10.1186/s13059-019-1910-1

76. Pertea, G., & Pertea, M. (2020). GFF utilities: GffRead and GffCompare. F1000Research, 9, 304. 10.12688/f1000research.23297.2

77. Stanke, M., Diekhans, M., Baertsch, R., & Haussler, D. (2008). Using native and syntenically mapped cDNA alignments to improve de novo gene finding. Bioinformatics (Oxford, England), 24(5), 637–644. 10.1093/bioinformatics/btn013

78. Stanke, M., Schöffmann, O., Morgenstern, B., & Waack, S. (2006). Gene prediction in eukaryotes with a generalized hidden Markov model that uses hints from external sources. BMC Bioinformatics, 7(1), 62. 10.1186/1471-2105-7-62

79. Gabriel, L., Hoff, K. J., Brůna, T., Borodovsky, M., & Stanke, M. (2021). TSEBRA: transcript selector for BRAKER. BMC Bioinformatics, 22(1), 566. 10.1186/s12859-021-04482-0

80. Chan, P. P., Lin, B. Y., Mak, A. J., & Lowe, T. M. (2021). tRNAscan-SE 2.0: improved detection and functional classification of transfer RNA genes. Nucleic Acids Research, 49(16), 9077–9096. 10.1093/nar/gkab688

81. Dainat, J., Hereñú, D., Murray, K. D., Davis, E., Ugrin, I., Crouch, K., et al. (2024). Another GFF/GTF Analysis Toolkit. Retrieved from https://zenodo.org/records/13799920

82. Gremme, G., Steinbiss, S., & Kurtz, S. (2013). GenomeTools: a comprehensive software library for efficient processing of structured genome annotations. IEEE/ACM transactions on computational biology and bioinformatics, 10(3), 645–656. 10.1109/TCBB.2013.68

83. Jones, P., Binns, D., Chang, H.-Y., Fraser, M., Li, W., McAnulla, C. et al. (2014). InterProScan 5: genome-scale protein function classification. *Bioinformatics (Oxford*, England), 30(9), 1236–1240. 10.1093/bioinformatics/btu031

84. Huerta-Cepas, J., Szklarczyk, D., Heller, D., Hernández-Plaza, A., Forslund, S. K., Cook, H. et al. (2019). eggNOG 5.0: a hierarchical, functionally and phylogenetically annotated orthology resource based on 5090 organisms and 2502 viruses. Nucleic Acids Research, 47(D1), D309–D314. 10.1093/nar/gky1085

85. Cantalapiedra, C. P., Hernández-Plaza, A., Letunic, I., Bork, P., & Huerta-Cepas, J. (2021). EggNOG-mapper v2: Functional annotation, orthology assignments, and domain prediction at the metagenomic scale. Molecular Biology and Evolution, 38(12), 5825–5829. 10.1093/molbev/msab293

86. Käll, L., Krogh, A., & Sonnhammer, E. L. L. (2004). A combined transmembrane topology and signal peptide prediction method. Journal of Molecular Biology, 338(5), 1027–1036. 10.1016/j.jmb.2004.03.016

87. Sharaf, A., & Voolstra, C. R. (2025). SequAna-Ukon/GeneForge: GeneForge v1.0. Zenodo. 10.5281/ZENODO.16631467

88. Hoff, K. J., Lomsadze, A., Borodovsky, M., & Stanke, M. (2019). Whole-genome annotation with BRAKER. *Methods in Molecular Biology (Clifton*, N.J.), 1962, 65–95. 10.1007/978-1-4939-9173-0_5

89. Fiesinger, A., Buitrago-López, C., Sharaf, A., Cárdenas, A., & Voolstra, C. (2024). A draft genome assembly of the reef-building coral Acropora hemprichii from the central Red Sea. Scientific Data, 11(1), 1288. 10.1038/s41597-024-04080-8

90. Shinzato, C., Khalturin, K., Inoue, J., Zayasu, Y., Kanda, M., Kawamitsu, M. et al. (2021). Eighteen coral genomes reveal the evolutionary origin of Acropora strategies to accommodate environmental changes. Molecular Biology and Evolution, 38(1), 16–30. 10.1093/molbev/msaa216

91. Buitrago-López, C., Mariappan, K. G., Cárdenas, A., Gegner, H. M., & Voolstra, C. R. (2020). The Genome of the Cauliflower Coral Pocillopora verrucosa. Genome Biology and Evolution, 12(10), 1911–1917. 10.1093/gbe/evaa184

92. Pu, Y., Zhou, Y., Liu, J., & Zhang, H. (2024). A high-quality chromosomal genome assembly of the sea cucumber Chiridota heheva and its hydrothermal adaptation. GigaScience, 13. 10.1093/gigascience/giad107

93. Liang, Y., Xu, K., Li, J., Shi, J., Wei, J., Zheng, X. et al. (2025). The molecular basis of octocoral calcification revealed by genome and skeletal proteome analyses. GigaScience, 14. 10.1093/gigascience/giaf031

94. Lu, Y., Luo, F., Zhou, A., Yi, C., Chen, H., Li, J. et al. (2024). Whole-genome sequencing of the invasive golden apple snail Pomacea canaliculata from Asia reveals rapid expansion and adaptive evolution. GigaScience, 13. 10.1093/gigascience/giae064

95. Wang, Y.-S., Li, M.-Y., Li, Y.-L., Li, Y.-Q., Xue, D.-X., & Liu, J.-X. (2024). Chromosome-level genome assemblies of two littorinid marine snails indicate genetic basis of intertidal adaptation and ancient karyotype evolved from bilaterian ancestors. GigaScience, 13. 10.1093/gigascience/giae072

96. Arantes, L. S., Brown, T., De Panis, D., Whiting, S. D., Young, E. J., LaCasella, E. L., et al. (2025). Haplotype-resolved reference genomes of the sea turtle clade unveil ultra-syntenic genomes with hotspots of divergence. GigaScience, 14(giaf105). 10.1093/gigascience/giaf105

